# Reduced male fertility of an Antarctic mite following extreme heat stress could prompt localized population declines

**DOI:** 10.1101/2023.04.11.535735

**Authors:** Joshua B. Benoit, Geoffrey Finch, Andrea L. Ankrum, Jennifer Niemantsverdriet, Bidisha Paul, Melissa Kelley, J. D. Gantz, Stephen F. Matter, Richard E. Lee, David L. Denlinger

**Affiliations:** Department of Biological Sciences, University of Cincinnati, Cincinnati, OH, USA; Department of Environmental Health, University of Cincinnati, Cincinnati, OH, USA; Department of Biology, Miami University, Oxford, OH, USA; Department of Biology and Health Science, Hendrix College, Conway, AR, USA; Departments of Entomology and Evolution, Ecology and Organismal Biology, The Ohio State University, Columbus, OH, USA

**Keywords:** Antarctic mite, *Alaskozetes antarcticus*, reproduction, climate change, heat stress

## Abstract

Climate change is leading to substantial global thermal changes, which are particularly pronounced in polar regions. Few studies have examined the impact of heat stress on reproduction in Antarctic terrestrial arthropods, specifically how brief, extreme events may alter survival. We observed that sublethal heat stress reduces male fecundity in an Antarctic mite, yielding females that produced fewer viable eggs. Females and males collected from microhabitats with high temperatures showed a similar reduction in fertility. This impact is temporary, as indicated by recovery of male fecundity following return to cooler, stable conditions. The diminished fecundity is likely due to a drastic reduction in the expression of male-associated factors that occur in tandem with a substantial increase in the expression of heat shock proteins. Cross mating between mites from different sites confirmed that heat-exposed populations have impaired male fertility. However, the impact on fertility declines with time when the mites are allowed to recover under less stressful conditions, suggesting that the negative effects are transient. Modeling indicated that heat stress is likely to reduce population growth and that short bouts of non-lethal heat stress could have substantial effects on local populations of Antarctic arthropods.

## Introduction

The climate changes occurring on Earth are causing many populations to decline or become extinct (Thomas et al. 2004; Keller et al. 2008; Chen et al. 2011), and these effects are most rapid in polar regions (Turner and Marshall 2011), where changes in the terrestrial environment are the most drastic. Terrestrial animals in Antarctica are limited to invertebrates, and one of the terrestrial invertebrates, the Antarctic oribatid mite, *Alaskozetes antarcticus*, is among the most common. This mite is found in large aggregations with hundreds to thousands of individuals, presenting in all developmental stages (Block and Convey 1995). Sperm are transferred to the females externally; males deposit stalked spermatophores that fertilize the females through her genital aperture (Block and Convey 1995). A single molt occurs each year, and 4-5 years are required for the mite to reach maturation (Convey 1994a; Marshall and Convey 1999; van Vuuren et al. 2018)(van Vuuren et al. 2018).

This mite has been the focus of numerous studies on environmental stress tolerance that have monitored survival and metabolism (Young and Block 1980; Hayward et al. 2003; Benoit et al. 2008; Everatt et al. 2013). Yet, little is known about the more specific effects of thermal change on mite physiology. Importantly, the temperatures at lethal or viability thresholds (critical thermal limit, CTL) often overestimate survival and proliferation of populations. Specifically, male and female fertility can be greatly impacted by thermal changes, resulting in populations becoming non-viable after exposure to short periods of high temperature (Krebs and Loeschcke 1994; David et al. 2005; Zizzari et al. 2011; Sales et al. 2018; Walsh et al. 2019; Iossa 2019; van Heerwaarden and Sgrò 2021). The thermal fertility limit (TFL) of a species represents an overlooked aspect where sublethal temperature may impact species’ persistence (Walsh et al. 2019; van Heerwaarden and Sgrò 2021). Importantly, tropical species tend to have a higher temperature limit of fertility than temperate species (Rohmer et al. 2004). These fertility-associated effects can even be transgenerational (Sales et al. 2018), with reduced survival and reproductive potential in surviving offspring. Few studies have examined the impact of thermal exposure on the fertility of polar invertebrates. Factors that affect species population persistence at sublethal levels need to be assessed, as thermal bouts are expected to be more frequent and more extended as climate change progresses (Meehl and Tebaldi 2004).

In this study, we examined the effect of short bouts of ecologically relevant heat stress on the fertility of male Antarctic mites. We observed that mites from warm microhabitats produce substantially fewer viable eggs. We noted a reduction in fertility when males from heat-stressed areas were mated with females from sites not exposed to heat stress. RNA-seq reveals that male thermal stress leads to increased expression of heat shock proteins and a general reduction in male-enriched transcripts. Short heat bouts could substantially reduce population growth for polar invertebrates and greatly impact the genetic diversity of polar mite populations that inherently have a limited distribution and lack dispersal options.

## Methods

### Mite collections

Mites were collected from three islands (Humble, Christine, and Cormorant) near Palmer Station, Antarctica. Thermal changes were monitored at each site using a HOBO (Onset Corp.) thermal logger for three days prior to the collection of the mites in January 2017. The mites were then held at 4 °C under long day length (20-h light: 4-h dark), conditions of summer at Palmer Station. Algae (*Prasiola crispa*) and other organic debris were provided. Sexes and juvenile stages (specifically tritonymphs), were separated based on previously established morphological characteristics (Block and Convey 1995; Meibers et al. 2019). A subset of tritonymphs were held individually to ensure emergence of virgin males and females.

### Egg viability from field collected sites and after treatments

Total egg counts were determined by dissection of a subset of females, which allowed for a census of developing eggs. For each island, egg viability was determined by collecting eggs from females that were placed in a small dish with access to two males from the same island for one week. Food sources were replaced, and *P. crispa* and other organic debris were examined for the presence of eggs. The number of juvenile mites was tabulated as a proxy for egg viability at 50, 100 and 150 days. These times were selected to encompass two summer equivalents of degree days, which is the noted time required for mites to deposit all eggs and to allow time for emergence. For heat-based studies and in cross mating of mites from different islands, tritonymphs were collected and stored with ample food sources. Every 12 h, females were removed and stored to provide control, virgin females. Following treatments, groups of three males were placed with a single female and juvenile emergence was assessed as before.

Differences between total eggs produced and egg viability levels were compared with ANOVA followed by a Tukey’s test.

### Thermal stress exposure of male mites

Male mites were exposed to thermal stress based on previous studies (Rosendale et al. 2016a; Fieler et al. 2021). Groups of 8 mites were held in 1.5 ml tubes and placed within foam plugged 50 mL centrifuge tubes. One of these centrifuge tubes was filled with a supersaturated solution of potassium nitrate to maintain a relative humidity of 93%. The 50 mL tubes were suspended in an ethylene glycol:water (60:40) solution. The temperature was controlled with a programmable cold bath (Arctic A25; Thermo Scientific, Pittsburgh, PA, USA) set at 30 °C to mimic the extreme thermal stress observed under field observations (Fig. 1). Following treatments, mites were returned to colony conditions and allowed to recover for 24 h prior to survival assessment. After 6 h, a second subset of males was transferred directly to -70°C for RNA extraction. Mites were marked as surviving if the larvae could move five body lengths following probing. Lastly, a final subset of males was used in mating studies.

**Figure 1:**
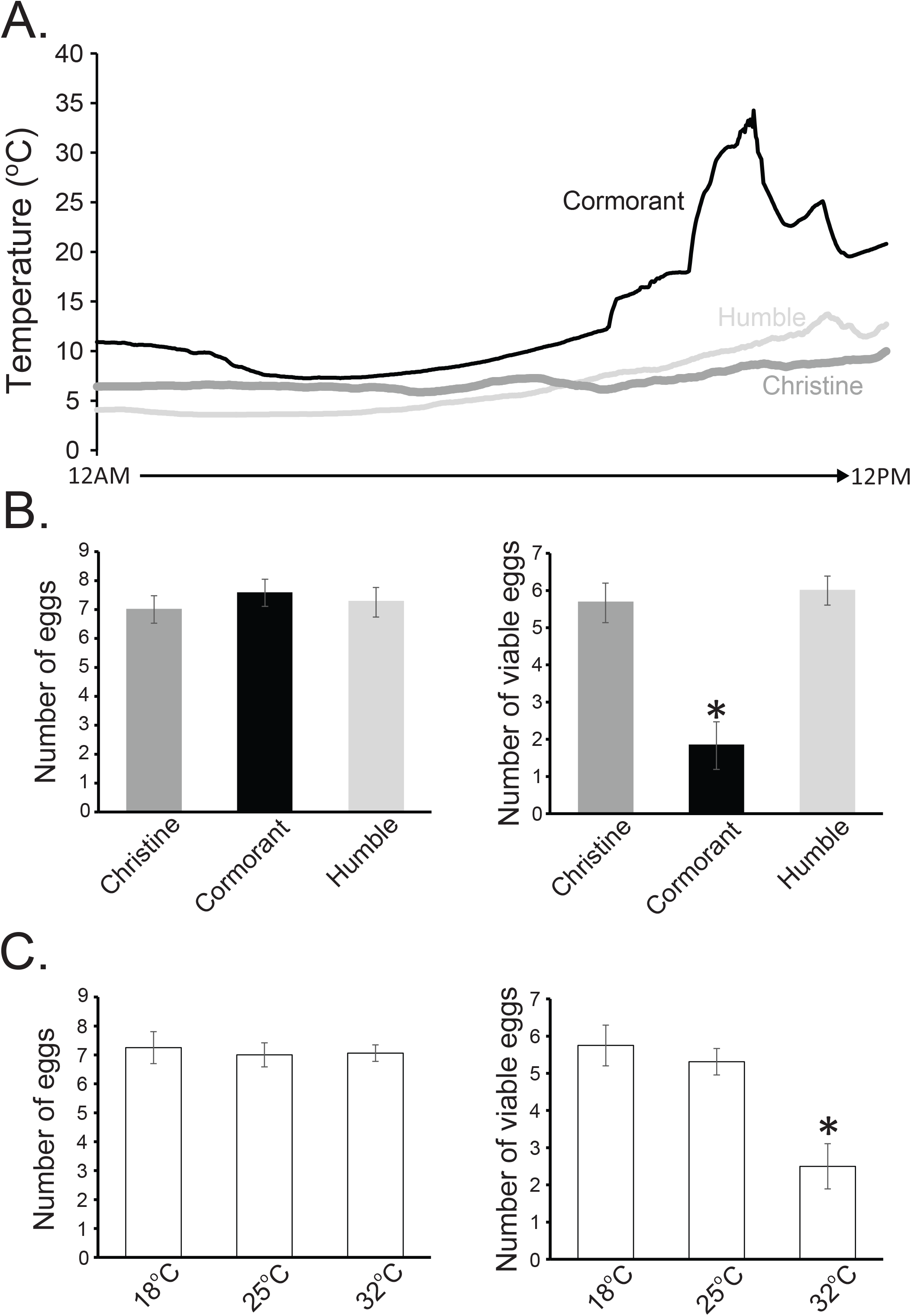
Thermal stress directly impacts Antarctic mite egg production. A. Thermal profile of three island sites used for collection of Antarctic mites, *Alaskozetes antarcticus*, near Palmer Station, Antarctica. B. Total developing eggs observed within females by dissection (left) and viable egg number based on the number of observed nymphs (right). C. Impact of male thermal stress on egg viability for mites collected from Christine Island. Total developing eggs observed within females by dissection (left) and viable egg number based on the number of observed nymphs (right). *, indicates significantly different at 0.05 based on ANOVA. Each sample was based on 10-12 replicates.

### RNA-seq analyses following heat stress

RNA was extracted from groups of 20 mites by homogenization (BeadBlaster 24, Benchmark Scientific) in Trizol (Invitrogen) based on other acarine studies (Meibers et al. 2019; Rosendale et al. 2022). DNase I (Thermo Scientific) was used to remove contaminating DNA and each sample was cleaned with a GeneJet RNA Cleanup and Concentration Micro Kit (Thermo Scientific) according to the manufacturer’s protocols. RNA concentration and quality were examined with a NanoDrop 2000 (Thermo Scientific) followed by an Agilent Bioanalyzer. Three independent biological replicates were generated for the control and heat-stressed males.

Samples were processed and sequenced at the DNA Sequencing and Genotyping Core at Cincinnati Children’s Hospital and Medical Center. A Qubit 3.0 Fluorometer (Life Technologies) was used to quantify and allow for 200 ng of RNA to be used for each sample. RNA was poly(A) selected and reverse transcribed using a TruSeq Stranded mRNA Library Preparation Kit (Illumina). Each sample was barcoded (8-base) for multiplex analysis and amplified (15 cycle PCR). Each library was sequenced on a HiSeq 2500 sequencing system (Illumina) in Rapid Mode. Each sample was sequenced with a minimum depth of 30 million paired-end reads (75 bp). RNA-seq data have been deposited at the National Center for Biotechnology Information (NCBI) Sequence Read Archive (Bioproject: PRJNA951800).

A previously published *de novo* assembly based on sex-specifc analyses was used in the RNA-seq analysis to allow for direct comparison to male-enriched gene sets (Meibers et al. 2019). Two separate pipelines were used to assess differential transcript expression. The first method was with CLC Genomics (Qiagen, version 10) based on previous methods (Benoit et al. 2014; Rosendale et al. 2016b). Reads mapped to the contigs were at least 85% of the reads matching at 90% with a mismatch allowed of two. Each read was allowed to align to 20 contigs.

A Baggerly’s test (a proportion-based statistical test) was used to test significance among samples. Multiple comparison correction was performed (false discovery rate, FDR). The second method utilized was Kallisto (Bray et al. 2016) under default setting. Differential expression between contigs was examined with the DESeq2 package (Love et al. 2014). A generalized linear model assuming a binomial distribution followed by FDR approach was used to account for multiple tests (Benjamini and Hochberg 1995). Cut-off values for significance were determined as described in the CLC-based analyses. After identification of the contigs with differential expression, enriched functional groups were identified with gProfiler (Reimand et al. 2018).

### Quantitative PCR of targeted male-enriched genes

Expression levels for targeted genes were measured based on previous methods (Rosendale et al. 2016b; Finch et al. 2020). RNA was extracted as described above. Complementary DNA (cDNA) was generated with a DyNAmo cDNA Synthesis Kit (Thermo Scientific). Each reaction used 100 ng RNA, 50 ng oligo (dT) primers, the reaction buffer containing dNTPs, 5 mM MgCl_2_, and M-MuLV RNase H+ reverse transcriptase. KiCqStart SYBR Green qPCR ReadyMix (Sigma Aldrich), along with 300 nM forward and reverse primers, cDNA diluted 1:20, and nuclease–free water were used for all reactions. Primers used were designed with Primer3 (Kõressaar et al. 2018). All qPCR reactions were conducted using an Illumina Eco quantitative PCR system. Four biological replicates were examined for each gene. Expression levels were normalized to alpha-tubulin using the ΔΔCt methods. Differences between expression levels were compared with ANOVA followed by a Tukey’s test.

### Population level effects

To assess the impact of a short heat exposure on mite populations, we used a Leslie matrix approach (Lefkovitch 1965) previously used to assess population changes for other Antarctic terrestrial arthropods (Finch et al. 2020). The dominant eigenvalue for the matrix is the population growth rate (λ). The life history was simplified into three stages: egg, pre-reproductive, and reproductive. We assumed no within stage transition due to a lack of within stage survivorship data, and based estimates on development and reproduction occurring over the course of five years (Convey 1994b; Block and Convey 1995). For mites under stable conditions (control), a mean fecundity of 7 eggs with a viability of 0.85 was assumed. For all groups we assumed a 0.90 probability of transition from pre-reproductive to reproductive and mortality after. For mite populations that experienced a single thermal bout, egg viability declined to 0.55 while for those that experienced two thermal bouts, egg viability declined to 0.25. For these analyses, a single fertilization event was assumed to occur, rather than multiple, as the impact of multiple mating events has not been studied. The dominant eigenvalue of each matrix was determined using the function “eigen” in R.

## Results

### Differences in egg viability from mites collected at different sites

When we collected the mites we noted distinct thermal differences at the collecting sites (Fig. 1A). Specifically, temperatures on Cormorant Island were considerably higher than at the other sites. Mites collected near the sites of the temperature probes deposited the same number of eggs (ANOVA, P > 0.1 for all cases) and different levels of egg viability based on the number of observed progeny (ANOVA, d.f._2,33_, F = 32.21, P < 0.001). Egg viability was reduced in females collected from Cormorant Island (Tukey’s, P < 0.0001; Fig. 1B). To determine if a rapid bout of heat stress is the likely reason for the decline in fertility, male mites were collected from Humble and exposed to thermal stress to mimic those observed in the population collected from Cormorant Island (Fig. 1C, ANOVA, d.f._2,45_, F = 21.16, P < 0.001). Importantly, during collection, we observed numerous male mites moving on the warm surface, while most females were protected underneath rocks and other debris, suggesting the males are more likely to be exposed to extreme thermal differences.

### RNA-seq analyses revealed thermal stress is likely to impact male-associated factors

When male mites were exposed to heat stress, there were only 46 contigs with increased expression (Fig. 2A, FDR < 0.05, Table S1). These genes were predominantly associated with response to heat stress, including a multitude of canonical heat shock proteins (Fig. 2B). Heat stress resulted in suppression of nearly 800 contigs, including a significant enrichment for contigs previously identified as male-associated (Fig. 2A,C, Table S2). Of these suppressed genes, there was enrichment for GO categories associated with serine/threonine activity, including many contigs that are likely testis-specific (Fig. 2C,D). This transcriptional profile highlights the observation that male-associated factors, specifically those that underlie testis function, declined in abundance. Results between specific RNA-seq pipelines gave considerable overlap (Pearson correlation = 0.913) with nearly identical functional GO categories, thus we only reported the RNA-seq analysis pipeline with Kallisto mapping and DESeq2 for statistical analyses. The general transcriptome results highlight that a short thermal bout increases the expression of stress-associated genes at the expense of the male-associated factors.

**Figure 2:**
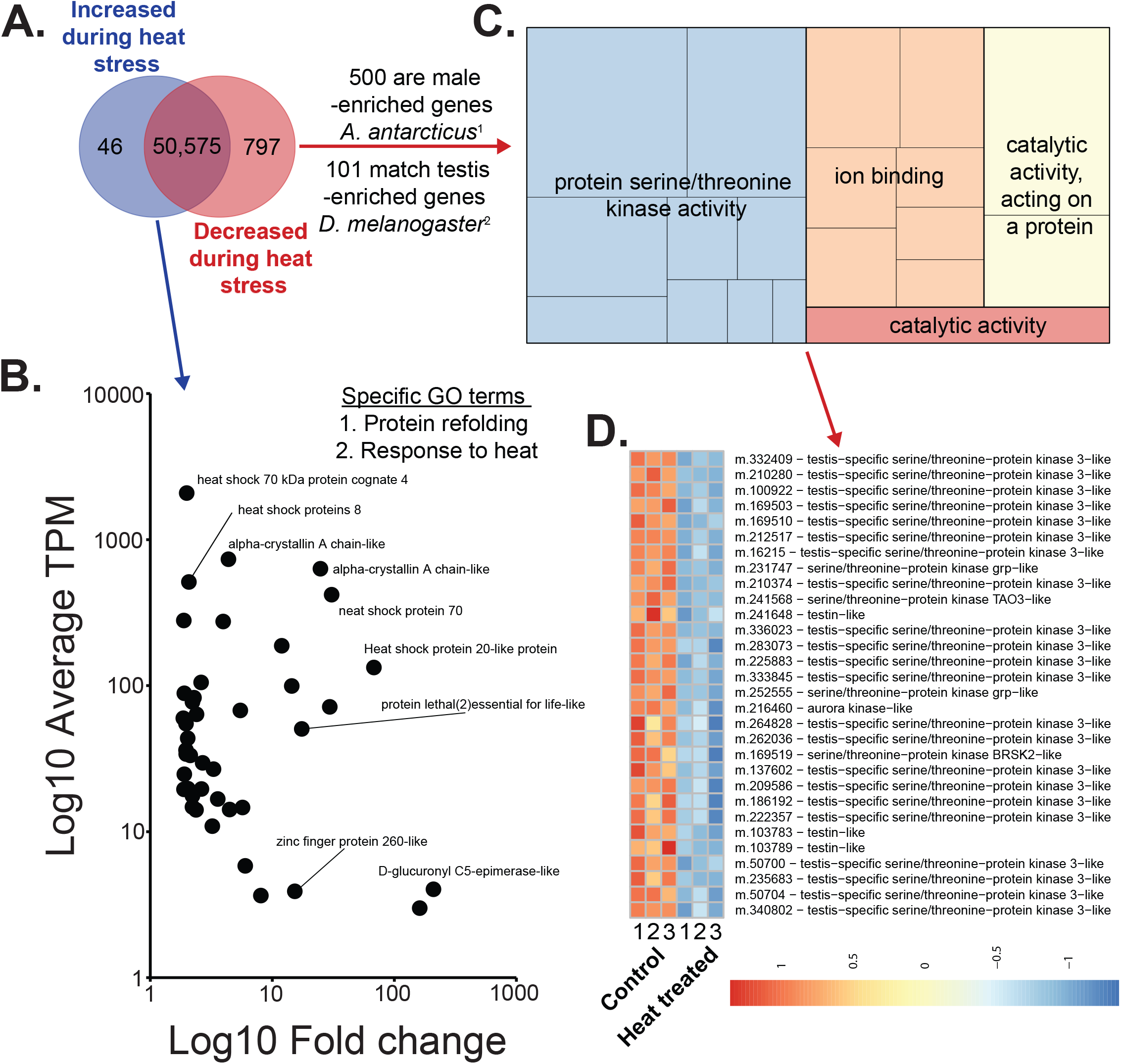
RNA-seq analyses reveal distinct male-associated changes following heat stress in the Antarctic mite. A. Total contigs with differential expression following exposure to heat stress. A significant number of these are male-enriched based on a comparison to a previous sex-specific analysis (Meibers et al. 2019) and on a Fisher’s exact test (P < 0.05). Contig assembly from Meibers et al. (Meibers et al. 2019). B. Contigs with increased expression are associated with the heat shock response. TPM, transcript per million. C. Gene ontology (GO) for contigs with decreased expression following heat treatment. Color areas represent higher order GO categories with smaller blocks presenting lower order that are contained within the color GO. D. Expression patterns of contigs that likely underlie sperm/testis development, all showing decreased expression following heat exposure.

**Table 1.** Modeled population growth of *A. antarcticus* under heat stress conditions indicates that multiple heat shock exposures will decrease population growth.

| Treatment | Year | Times | Population growth ($\lambda$) |
| --- | --- | --- | --- |
| Control | 1 | 0 | 1.75 |
| Heat Shock | 1 | 1 | 1.51 |
| Heat Shock | 1 | 2 | 0.58 |

### Expression levels of specific genes vary for mites collected on different islands

To confirm whether males from Cormorant Island experienced heat stress, we measured the expression levels for six contigs identified in the RNA-seq studies to be responsive to heat stress (Fig. 3). Four of the male-enriched contigs showed significantly decreased expression in male mites collected from Cormorant Island compared to those from Humble and Christine Islands (Fig. 3A-D, ANOVA P < 0.05). By contrast, two proteins associated with the heat shock response were elevated in mites from Cormorant (Fig. 3E,F, ANOVA P < 0.05). These studies confirm that male mites from Cormorant, but not Christine or Humble, are experiencing heat stress that is likely impacting aspects of male fertility identified in the RNA-seq studies.

**Figure 3:**
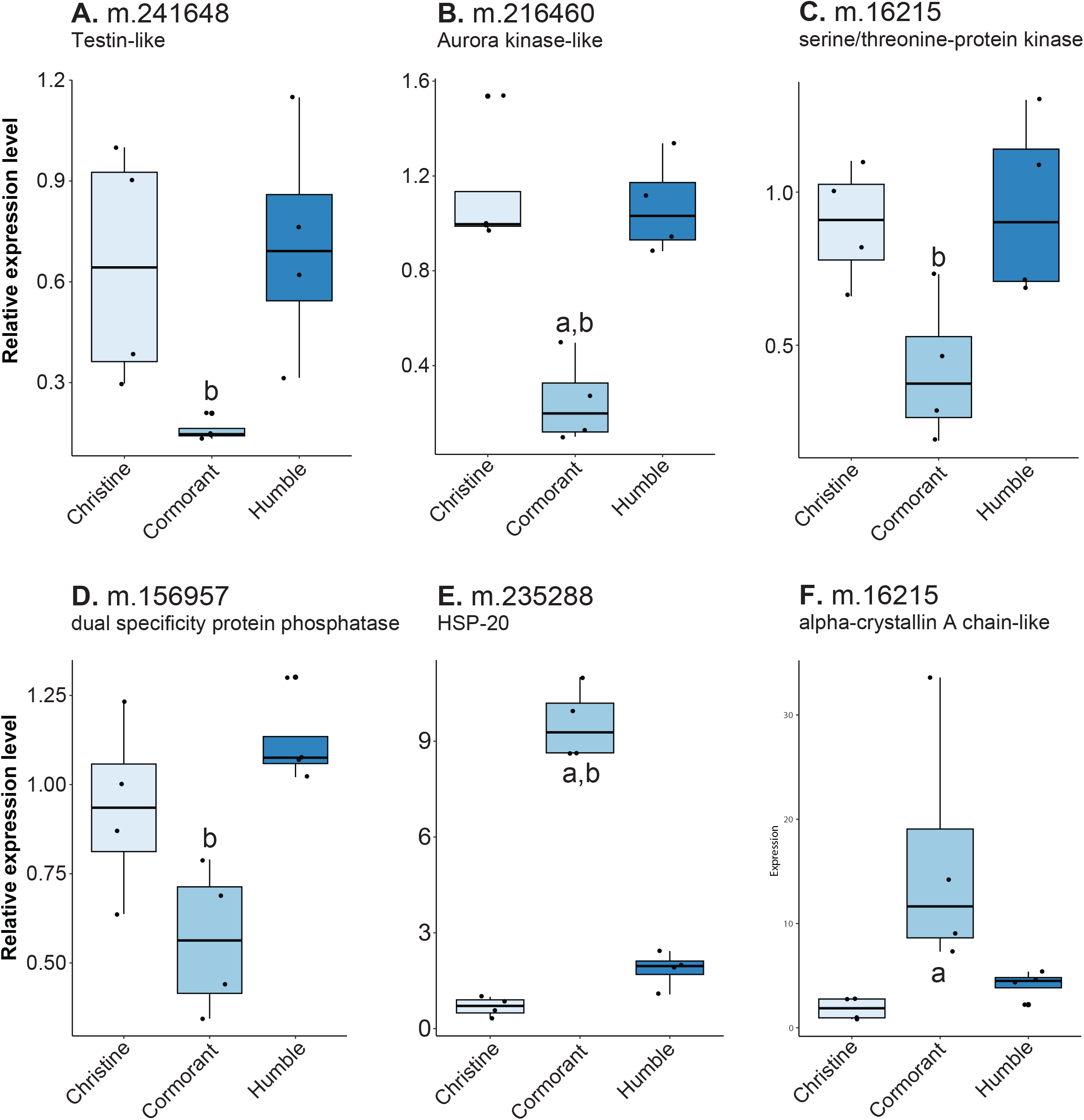
Male-enriched contigs have reduced transcript levels in Antarctic mites collected from Cormorant Island. A-D. Male-enriched contigs. E-F. Heat stress associated contigs. a, significant when compared to mites collected from Christine Island. b, significant when compared to mites collected from Humble Island. qPCR was based on four replicates. Significance was determined with ANOVA followed by a Tukey test.

### Reduced fertility leads to potential declines in population levels

When males from different islands were cross-mated, there were significant declines in fertility (Fig. 4A-B, ANOVA, d.f._3,44_, F = 17.62, P < 0.001). Specifically, female mites fertilized by males from Cormorant Island had reduced fecundity (Fig. 4A, Tukey’s, P < 0.001). If males were allowed to recover for a period of two weeks, this reduction in fertility was eliminated, indicating the impact of heat stress is transient (Fig. 4A, Tukey’s, P < 0.001). In mites from Humble Island that were exposed to heat stress, a reduction in male fertility similar to that noted in mites directly obtained from Cormorant Island was observed (Fig. 4A, Tukey’s, P < 0.001). This reduction was also transient, with recovery occurring after two weeks under stable conditions (Fig. 4A). These studies confirm that brief periods of thermal stress are likely a major factor in the decline of male fertility.

**Figure 4:**
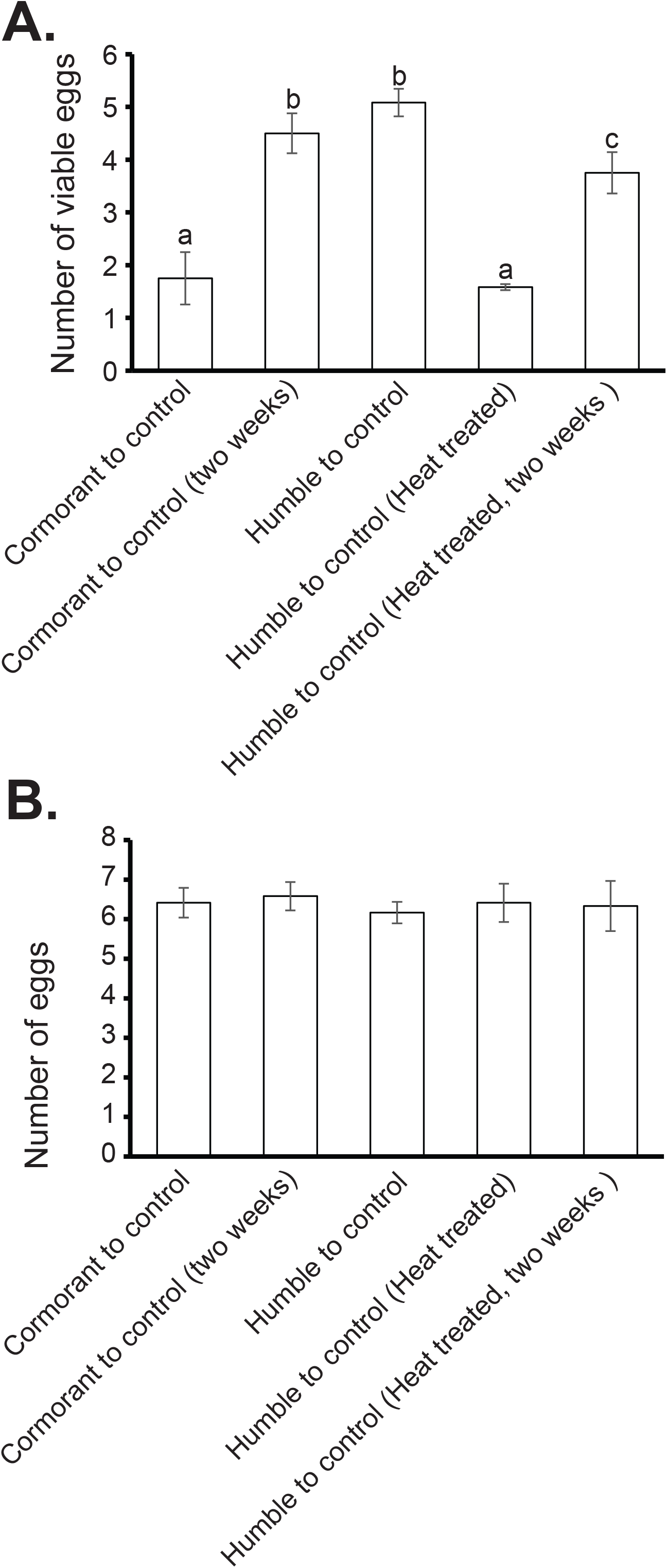
Mating of male mites to individuals from different islands to confirm heat stress reduces fecundity. (Top) Impact of male status on egg viability for mites. Total developing eggs observed within females by dissection (left) and viable egg number based on the number of observed nymphs (right). Each sample was based on 10-12 replicates. Significance was determined with ANOVA followed by a Tukey test.

Repeated bouts of heat stress and the associated reduction in egg survival are sufficiently strong to impact population growth. Under control conditions, population growth for mites was estimated as λ = 1.75 (where λ = 1 is the replacement rate). One bout of heat stress reduced population growth to λ = 1.51, but the population still showed positive growth. However, a second bout of heat stress reduced population growth to below replacement (λ = 0.58), resulting in a population decline.

## Discussion

Our studies provide direct evidence that male fertility in Antarctic mites will be impacted by high temperature exposure. This is likely to occur under natural conditions as male mites collected from areas of high temperature exposure show reduced fertility. Transcript level assessment indicated that males show increased expression of genes associated with repair and prevention of damage due to heat shock. This response seems to occur at the expense of factors that underlie male fertility, specifically those associated with testis function. Cross-mating between mites from different islands and direct heat exposure confirmed that short thermal stress bouts cause lower male fertility. Lastly, our population growth modeling suggests that these short heat bouts are likely to result in declines in local populations. Importantly, our studies were conducted on naturally-occurring populations and under thermal exposures already occurring, highlighting that these studies are likely to represent frequent and common thermal stresses.

Heat stress bouts have a multitude of effects on survival, growth, and reproduction of terrestrial arthropods (Sales et al. 2018, 2021; Denlinger and Yocum 2019; Walsh et al. 2019). Aspects related to fertility are altered by thermal stress and range from changes in courtship behavior to direct impacts on sperm function (Araripe et al. 2004; Walsh et al. 2019). Here, we observed that male mites have reduced fertility when collected from locations that recently experienced thermal stress. Importantly, as this mite produces external spermatophores, thermal stress could impact sperm after being deposited externally (Block and Convey 1995). This suggests that the decline in fertility could be the result of two different factors, a reduction in the production of external spermatophores and/or spermatophores that contain sperm of lower quality with impaired fertilization capability. In most other systems, the impact of male fertility was measured in animals with direct copulation (Araripe et al. 2004; David et al. 2005; Walsh et al. 2019), rather than the indirect process used by *A. antarcticus*. The indirect fertilization used by mites could result in dual consequences of thermal stress: males could be directly impacted by heat stress during sperm production and/or spermatophores could be affected by thermal exposure once placed in the environment. Our field studies show that egg viability could be affected by both processes, but cross-mating studies suggest that direct impacts on the male likely represent a major influence.

A major consequence of thermal stress in diverse systems is defects in sperm function (David et al. 2005; Vasudeva et al. 2014; Sales et al. 2018). Here, we did not directly observe sperm function, but instead we note that heat stress yielded a substantial decline in previously identified male-enriched factors (Meibers et al. 2019) likely required for both the generation of sperm and seminal fluid (Findlay et al. 2008; Avila et al. 2010; Scolari et al. 2016; Meibers et al. 2019). Specifically, heat stress results in a critical decline in serine/threonine kinase (TSSK), an enzyme linked to sperm viability (Spiridonov et al. 2005; Xu et al. 2008). Serine/threonine kinase transcripts are abundant in males or male reproductive organs of multiple mites (Sonenshine et al. 2011; Joag et al. 2016; Mondet et al. 2018). These expressional profile changes suggest that sperm generation is likely impaired or poor quality sperm are generated in heat-stressed *A. antarcticus*. When targeted genes were measured directly from field samples, mites from Cormorant Island had reduced expression levels of male-enriched factors and increased levels of heat shock genes. These studies suggest that thermal stress causes males to either generate fewer sperm or sperm of lower quality.

Studies on reproduction of arthropods in Antarctica are limited, with most studies providing a basic description of reproductive output, while only a few examined effects at the molecular level (Convey 1998). Two recent studies reported transcriptional changes in male and female reproduction; one study was used to identify male-associated factors in *A. antarcticus* (Meibers et al. 2019) and a second examined male-enriched aspects in the Antarctic midge, *Belgica antarctica* (Meibers et al. 2019; Finch et al. 2020). In addition, acclimation to thermal stress is known to increase fertility in the Antarctic midge (Ajayi et al. 2021). Critical thermal limits in relation to survival *of A. antarcticus* suggest that this mite can tolerate temperatures far above those experienced in these studies (Block and Convey 1995; Hayward et al. 2003; Everatt et al. 2013) but at a significant cost in fertility. By examining fertility limits, we showed that local mite populations are likely to decline rapidly due to fertility costs. Our modeling suggests that population declines are likely to occur under field conditions that currently prevail in certain Antarctic sites, and that estimating the ability of Antarctic species to survive warming and more fluctuating temperatures due to climate change will require assessing fertility shifts during thermal stress.

## Acknowledgements

This work was supported by the National Science Foundation Grant DEB-1654417 (partially) and United States Department of Agriculture 2018-67013 (partially) to J.B.B. for shared equipment use, National Science Foundation grant OPP-1341393 to D.L.D., and National Science Foundation grant OPP-1341385 to R.E.L. We thank the staff at Palmer Station, Antarctica for assistance in logistics, experiments, and making the field season a pleasant and productive experience.

## Supplemental Tables

Table S1 - Contigs with increased expression during heat stress exposure in Alaskozetes *antarcticus* males.

Table S2 - Contigs with deceased expression during heat stress exposure in Alaskozetes *antarcticus* males.

## References

Ajayi OM, Gantz JD, Finch G, et al (2021) Rapid stress hardening in the Antarctic midge improves male fertility by increasing courtship success and preventing decline of accessory gland proteins following cold exposure. J Exp Biol 224.: https://doi.org/10.1242/jeb.242506

Araripe LO, Klaczko LB, Moreteau B, David JR (2004) Male sterility thresholds in a tropical cosmopolitan drosophilid, Zaprionus indianus. Journal of Thermal Biology 29:73–80

Avila FW, Sirot LK, LaFlamme BA, et al (2010) Insect seminal fluid proteins: identification and function. https://doi.org/10.1146/annurev-ento-120709-144823

Benjamini Y, Hochberg Y (1995) Controlling the false discovery rate: a practical and powerful approach to multiple testing. Journal of the Royal Statistical Society: Series B (Methodological) 57:289–300

Benoit JB, Attardo GM, Michalkova V, et al (2014) A novel highly divergent protein family identified from a viviparous insect by RNA-seq analysis: a potential target for tsetse fly-specific abortifacients. PLoS Genet 10:e1003874

Benoit JB, Yoder JA, Lopez-Martinez G, et al (2008) Adaptations for the maintenance of water balance by three species of Antarctic mites. Polar Biology 31:539–547

Block W, Convey P (1995) The biology, life cycle and ecophysiology of the Antarctic mite Alaskozetes antarcticus. Journal of Zoology 236:431–449

Bray NL, Pimentel H, Melsted P, Pachter L (2016) Erratum: Near-optimal probabilistic RNA-seq quantification. Nat Biotechnol 34:888

Chen I-C, Hill JK, Ohlemüller R, et al (2011) Rapid range shifts of species associated with high levels of climate warming. Science 333:1024–1026

Convey P (1994a) Growth and survival strategy of the Antarctic mite Alaskozetes antarcticus.N Ecography 17:97–107

Convey P (1994b) Growth and survival strategy of the Antarctic mite Alaskozetes antarcticus. Ecography 17:97–107

Convey P (1998) Latitudinal variation in allocation to reproduction by the Antarctic oribatid mite, Alaskozetes antarcticus. Applied Soil Ecology 9:93–99

David JR, Araripe LO, Chakir M, et al (2005) Male sterility at extreme temperatures: a significant but neglected phenomenon for understanding Drosophila climatic adaptations. J Evol Biol 18:838–846

Denlinger DL, Yocum GD (2019) Physiology of heat sensitivity. Temperature Sensitivity in Insects and Application in Integrated Pest Management 7–53

Everatt MJ, Bale JS, Convey P, et al (2013) The effect of acclimation temperature on thermal activity thresholds in polar terrestrial invertebrates. J Insect Physiol 59:1057–1064

Fieler AM, Rosendale AJ, Farrow DW, et al (2021) Larval thermal characteristics of multiple ixodid ticks. Comparative Biochemistry and Physiology Part A: Molecular & Integrative Physiology 257:110939

Finch G, Nandyal S, Perretta C, et al (2020) Multi-level analysis of reproduction in an Antarctic midge identifies female and male accessory gland products that are altered by larval stress and impact progeny viability. Sci Rep 10:19791

Findlay GD, Yi X, Maccoss MJ, Swanson WJ (2008) Proteomics reveals novel Drosophila seminal fluid proteins transferred at mating. PLoS Biol 6:e178

Hayward SAL, Worland MR, Convey P, Bale JS (2003) Temperature preferences of the mite, Alaskozetes antarcticus, and the collembolan, Cryptopygus antarcticus from the maritime Antarctic. Physiological Entomology 28:114–121

Iossa G (2019) Sex-specific differences in thermal fertility limits. Trends Ecol. Evol. 34:490–492

Joag R, Stuglik M, Konczal M, et al (2016) Transcriptomics of intralocus sexual conflict: gene expression patterns in females change in response to selection on a male secondary sexual trait in the bulb mite. Genome Biol Evol 8:2351–2357

Keller K, Tol RSJ, Toth FL, Yohe GW (2008) Abrupt climate change near the poles. Climatic Change 91:1–4

Kõressaar T, Lepamets M, Kaplinski L, et al (2018) Primer3_masker: integrating masking of template sequence with primer design software. Bioinformatics 34:1937–1938

Krebs RA, Loeschcke V (1994) Effects of exposure to short-term heat stress on fitness components in Drosophila melanogaster. Journal of Evolutionary Biology 7:39–49

Lefkovitch LP (1965) The study of population growth in organisms grouped by stages. Biometrics 21:1–18

Love MI, Huber W, Anders S (2014) Moderated estimation of fold change and dispersion for RNA-seq data with DESeq2. Genome Biol 15:550

Marshall DJ, Convey P (1999) Compact aggregation and life-history strategy in a continental Antarctic mite. Ecology and Evolution of the Acari 557–567

Meehl GA, Tebaldi C (2004) More intense, more frequent, and longer lasting heat waves in the 21st century. Science 305:994–997

Meibers HE, Finch G, Gregg RT, et al (2019) Sex- and developmental-specific transcriptomic analyses of the Antarctic mite, Alaskozetes antarcticus, reveal transcriptional shifts underlying oribatid mite reproduction. Polar Biology 42:357–370

Mondet F, Rau A, Klopp C, et al (2018) Transcriptome profiling of the honeybee parasite Varroa destructor provides new biological insights into the mite adult life cycle. BMC Genomics 19

Reimand J, Kolde R, Arak T (2018) gProfileR: interface to the g:Profiler toolkit. R package version 0 6 7:

Rohmer C, David JR, Moreteau B, Joly D (2004) Heat induced male sterility in Drosophila melanogaster: adaptive genetic variations among geographic populations and role of the Y chromosome. Journal of Experimental Biology 207:2735–2743

Rosendale AJ, Farrow DW, Dunlevy ME, et al (2016a) Cold hardiness and influences of hibernaculum conditions on overwintering survival of American dog tick larvae. Ticks Tick Borne Dis 7:1155–1161

Rosendale AJ, Leonard RK, Patterson IW, et al (2022) Metabolomic and transcriptomic responses of ticks during recovery from cold shock reveal mechanisms of survival. J Exp Biol 225.: https://doi.org/10.1242/jeb.236497

Rosendale AJ, Romick-Rosendale LE, Watanabe M, et al (2016b) Mechanistic underpinnings of dehydration stress in the American dog tick revealed through RNA-Seq and metabolomics. J Exp Biol 219:1808–1819

Sales K, Vasudeva R, Dickinson ME, et al (2018) Experimental heatwaves compromise sperm function and cause transgenerational damage in a model insect. Nature Communications 9

Sales K, Vasudeva R, Gage MJG (2021) Fertility and mortality impacts of thermal stress from experimental heatwaves on different life stages and their recovery in a model insect. Royal Society Open Science 8

Scolari F, Benoit JB, Michalkova V, et al (2016) The spermatophore in Glossina morsitans morsitans: insights into male contributions to reproduction. Sci Rep 6:20334

Sonenshine DE, Bissinger BW, Egekwu N, et al (2011) First transcriptome of the testis-vas deferens-male accessory gland and proteome of the spermatophore from Dermacentor variabilis (Acari: Ixodidae). PLoS One 6:e24711

Spiridonov NA, Wong L, Zerfas PM, et al (2005) Identification and characterization of SSTK, a serine/threonine protein kinase essential for male fertility. Mol Cell Biol 25:4250–4261

Thomas CD, Cameron A, Green RE, et al (2004) Extinction risk from climate change. Nature 427:145–148

Turner J, Marshall GJ (2011) Climate change in the polar regions

van Heerwaarden B, Sgrò CM (2021) Male fertility thermal limits predict vulnerability to climate warming. Nat Commun 12:2214

van Vuuren BJ, Lee JE, Convey P, Chown SL (2018) Conservation implications of spatial genetic structure in two species of oribatid mites from the Antarctic Peninsula and the Scotia Arc. Antarctic Science 30:105–114

Vasudeva R, Deeming DC, Eady PE (2014) Developmental temperature affects the expression of ejaculatory traits and the outcome of sperm competition in Callosobruchus maculatus. J Evol Biol 27:1811–1818

Walsh BS, Parratt SR, Hoffmann AA, et al (2019) The impact of climate change on fertility. Trends Ecol Evol 34:249–259

Xu B, Hao Z, Jha KN, et al (2008) Targeted deletion of Tssk1 and 2 causes male infertility due to haploinsufficiency. Dev Biol 319:211–222

Young SR, Block W (1980) Some factors affecting metabolic rate in an Antarctic mite. Oikos 34:178

Zizzari ZV, Valentina Zizzari Z, Ellers J (2011) Effects of exposure to short-term heat stress on male reproductive fitness in a soil arthropod. Journal of Insect Physiology 57:421–426

